# Syndecan-1 and thrombomodulin are early biomarkers for development of endotheliopathy in trauma and hemorrhagic shock

**DOI:** 10.1101/2023.04.03.535494

**Authors:** Tiffani C. Chance, Michael A. Meledeo, Andrew P. Cap, Daniel N. Darlington, James A. Bynum, Xiaowu Wu

## Abstract

The causes of endotheliopathy are multifactorial and trauma dependent, and the temporal mechanistic link that it has with acute traumatic coagulopathy (ATC) has yet to be fully determined. Therefore, we sought to define early characteristics and markers of endotheliopathy in two rat models, a time-course of acute lethal hemorrhage shock and polytrauma with hemorrhagic shock, to answer the following questions: (1) how soon can elevated biomarkers of endotheliopathy be detected in non-survivable (decompensated) hemorrhagic shock; (2) does extended hemorrhage time and accumulated hemorrhage impact biomarker levels; and (3) does the addition of polytrauma contribute to the further elevation of said biomarkers? In this study, we delineated a significant, acute elevation in end plasma levels of syndecan-1, thrombomodulin, and heparan sulfate, whose shedding patterns are a function of time as well as total hemorrhage volume vs. the addition of polytrauma. Additionally, we found that correlation of syndecan-1 and thrombomodulin to lactate levels and prothrombin times at trauma end revealed a potential for these markers to acutely predict downstream consequences of these trauma indications. Our results are of great relevance to the continued effort towards the identification and characterization of vascular dysfunction for early interventions in combat casualty care.

## Introduction

Endothelial cells play an integral role in the maintenance of vascular tone and integrity, maintenance of blood fluidity, transportation of fluid, nutrients, blood vessel formation and repair, modulation of the inflammatory response, and regulation of hemostasis, which are all important processes during and following trauma and hemorrhage (1). Trauma-linked endothelium damage (endotheliopathy, or increased vascular permeability after trauma) contributes to the enhancement of acute traumatic coagulopathy (ATC), tissue edema and hypoxia, and inflammation, which exacerbates hypovolemic shock and multiple organ failure (1-3).

The causes of endotheliopathy are multifactorial, and the temporal mechanistic link that it has with ATC (how ATC contributes to endotheliopathy and vice versa) has yet to be fully determined (4). Following acute trauma and hemorrhagic shock, early responses to changes in shear stress associated with alterations in blood flow due to hemorrhage (mechanotransduction), and later responses due to coagulopathy and subsequent shock, can lead to the degradation and shedding of the endothelial glycocalyx that normally serves as a shield for the endothelium as well as a permeability barrier of the vasculature (1, 5). The loss of glycocalyx causes endothelium to be activated and damaged by direct exposure to blood flow as well as circulatory disruptors that are released in the blood stream in response to trauma and hemorrhage, which can further contribute to the worsening of ATC (1, 5). This results in the shedding of key markers associated with endotheliopathy, such as syndecan (SDC-1; a shed product of the glycocalyx), thrombomodulin (TM; shed thrombin cofactor found on the surface of endothelial cells), and heparan sulfate (HS; shed product of the glycocalyx).

Work defining the correlation between SDC-1 and HS levels to glycocalyx degradation has previously been performed in a hemorrhaged rat model after treatment with different resuscitation fluids (6). This work, however, was performed with samples taken only at baseline and two hours post resuscitation - ignoring any acute changes in vascular structure and integrity that may occur in hemorrhage and shock. Conversely, Johansson et al. evaluated independent predictors of mortality in traumatic endotheliopathy in a prospective observational study of 424 severely injured patients upon hospital admission, but they were not able to isolate the effects of hemorrhage apart from combined injuries for an apt comparison of the ways in which compounded trauma (and the associated activation in endothelial cells, cytokines, and coagulopathy) can further affect endothelium health (7). Therefore, we sought to define early characteristics and biomarkers of endotheliopathy in a rat model of either lethal hemorrhage shock or of sublethal polytrauma with hemorrhage. This study seeks to answer the following three questions: (1) how soon can elevated biomarkers of endotheliopathy be detected following a decompensated hemorrhagic shock; (2) are biomarker levels associated with extended hemorrhage time and accumulated hemorrhage; and (3) does the addition of polytrauma contribute to the further elevation of said biomarkers?

## Results

A total of 33 rats were used to complete the 65% lethal hemorrhage model. T1 comprised the two blood draws at 4 and 6 min (1.5 ml/each), representing the changes in response to accumulated hemorrhage to 20% and 32% EBV within 4 to 6 min. T2 comprised the three blood draws at 8, 10, and 13 min (1.5 ml/each at 8 and 10 min; and 1 ml at 13 min) to represent the changes in response to accumulated hemorrhage from 39% to 51% EBV within 8 to 13 min. T3 consisted of a blood draw at either 16 (n = 16) or 19 min (n = 17), with subsequent datasets pooled, to represent the changes in response to accumulated hemorrhage of 56% or 57% EBV at 16 or 19 min. For subsequent data analysis, however, the data collected from T3 at either 16 or 19 min were pooled for analysis. A total of 47 rats were used to complete the polytrauma and 40% hemorrhage model. The final mL of blood drawn during hemorrhage, designated Last Hem, was drawn at 25.07 min ± 7.404 min from trauma start.

### SDC-1 sheds early following hemorrhage, with increased hemorrhage activating shedding compared to compounded polytrauma

SDC-1 concentrations over accumulating lethal hemorrhage (Fig. 1A) were significantly elevated from T2 (8.0 ng/ mL ± 1.8 ng/ mL) compared to T1 (6.4 ng/ mL ± 1.8 ng/ mL), T3 (13.6 ng/ mL ± 3.4 ng/ mL) compared to T2, and T3 compared to T1 (p < 0.05, p < 0.0001, p < 0.0001, respectively). SDC-1 concentrations were also detected early and at a significantly higher concentration (p < 0.01; Fig. 1B) when comparing reference control rats (4.5 ng/ mL ± 1.2 ng/ mL) to T1 blood draw (4 and 6 min). Both final blood draws, T3 and Last Hem (8.6 ng/ mL ± 1.6 ng/ mL), displayed significantly higher SDC-1 concentrations compared to control (Fig. 1C; p < 0.0001 and p < 0.001, respectively). Despite Last Hem (polytrauma and 40% hemorrhage) being pulled later than T3 (65% lethal hemorrhage shock), the higher combined final hemorrhage in T3 compared to Last Hem resulted in a significantly higher plasma concentration of SDC-1 (Fig. 1C; p < 0.0001).

**Figure 1.**
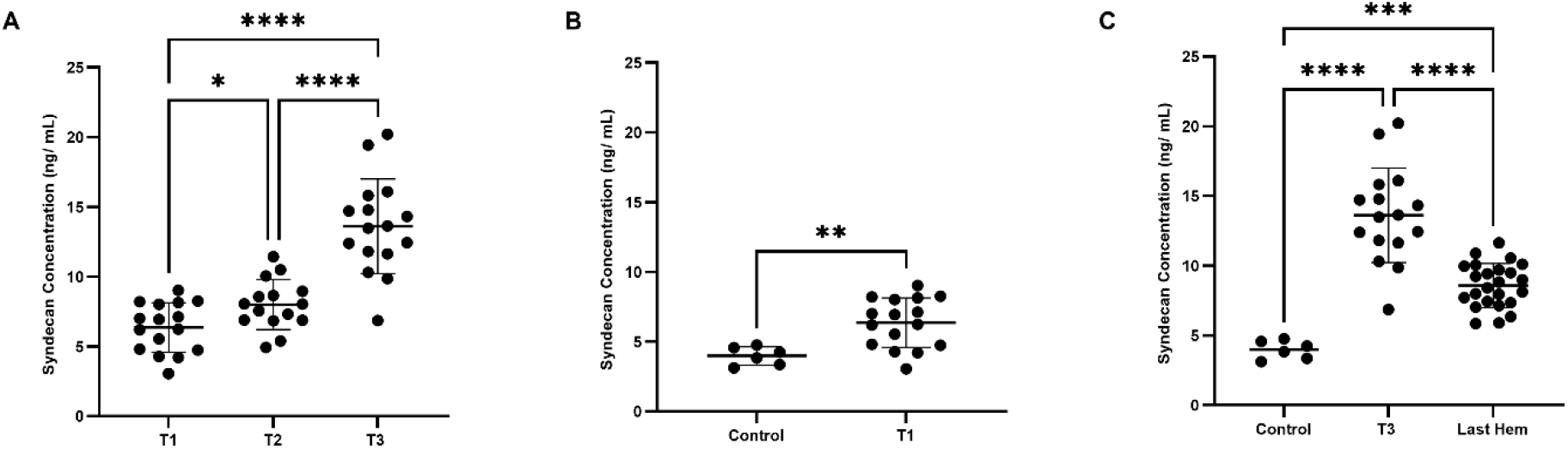
SDC-1 shedding in a rat model of lethal hemorrhage shock (65%) and polytrauma plus hemorrhage (40%). All data sets are displayed with individual data points, means, and standard deviations. Control denotes rats who underwent cannulation without trauma, and the blood sample was collected immediately followed by cannulation. (Fig. 1A) Repeated measures one-way ANOVA analysis of SDC-1 levels during accumulating time points. (Fig. 1B) Unpaired t-test analysis of Control and T1 plasma SDC-1 levels. (Fig. 1C) One-way ANOVA analysis Control, T3, and Last Hem plasma SDC-1 levels. * p ≤ 0.05, ** p ≤ 0.01, *** p ≤ 0.001, **** p ≤ 0.0001. “Last Hem” denotes last 1 mL of blood in rat with polytrauma and hemorrhage. T designates blood collected at various timepoints during our lethal hemorrhage shock model (see methods and Supplemental Figure 1). n = 16 for T1, n = 15 for T2, n = 16 for T3, n = 6 for control, and n = 24 for Last Hem.

### TM sheds early following hemorrhage, with increased hemorrhage causing no activation compared to compounded polytrauma

TM concentrations over accumulating lethal hemorrhage (Fig. 2A) were significantly elevated from T2 (3,324 pg/ mL ± 673pg/ mL) compared to T1 (2,598 pg/ mL ± 510pg/ mL), T3 (4,967 pg/ mL ± 1,345 pg/ mL) compared to T2, and T3 compared to T1 (p < 0.0001 for all). TM concentrations were also detected early and at a significantly higher concentration (p < 0.0001; Fig. 2B) when comparing reference control rats (1,171 pg/ mL ± 246pg/ mL) to T1 blood draw. Both final blood draws, T3 and Last Hem (5,090 pg/ mL ± 1,422 pg/ mL), displayed significantly higher TM concentrations compared to control (Fig. 1C; p < 0.0001 and p < 0.001, respectively).

**Figure 2.**
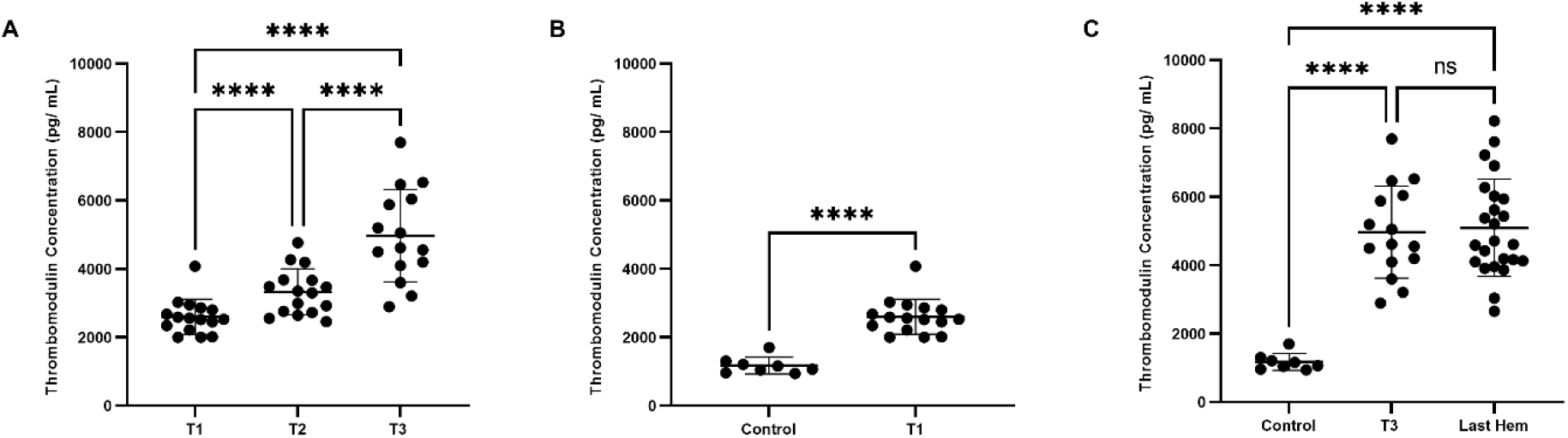
TM shedding in a rat model of lethal hemorrhage shock (65%) and polytrauma plus hemorrhage (40%). All data sets are displayed with individual data points, means, and standard deviations. Control denotes rats who underwent cannulation without trauma. (Fig. 2A) Repeated measures one-way ANOVA analysis of TM levels during accumulating time points denoted T1 (n = 16), T2 (n = 16), T3 (n = 15). (Fig. 2B) Unpaired t-test analysis of Control (and T1 plasma TM levels. (Fig. 2C) One-way ANOVA analysis Control (n = 8), T3, and Last Hem (n = 24) plasma TM levels. **** p ≤ 0.0001. Last Hem denotes plasma collected from the last 1 mL of blood of our rat polytrauma and hemorrhage model. T designates blood collected at various timepoints during our lethal hemorrhage shock model (see methods and Supplemental Figure 1). n = 16 for T1, n = 16 for T2, n = 15 for T3, n = 6 for control, and n = 37 for Last Hem.

Neither time of final hemorrhage blood draw between models nor final hemorrhage volume or the presence of additional trauma significantly impacted end TM plasma levels (Fig. 2C; p > 0.05).

### HS does not increase shedding with accumulating hemorrhage, time, nor compounded factor of polytrauma

HS concentrations over accumulating lethal hemorrhage (Fig. 3A) showed no significant changes (p > 0.05) between T1 (694.3 pg/ mL ± 478.6 pg/ mL), T2 (741.5 pg/ mL ± 440.8 pg/ mL), and T3 (958.1 pg/ mL ± 612.9 pg/ mL). HS concentrations were also not detected early and at a significantly higher concentration (p > 0.05; Fig. 3B) when comparing reference control rats (194.3 pg/ mL ± 16.57 pg/ mL) to T1 blood draw (4 and 6 min). Both final blood draws, T3 and Last Hem (1,387 pg/ mL ± 979.6 pg/ mL), displayed significantly higher HS concentrations compared to control (Fig. 3C; p < 0.05 for both). Time of final hemorrhage blood draw between models, nor final hemorrhage volume or presence of additional trauma, did not significantly impact end HS plasma levels (Fig. 3C; p > 0.05).

**Figure 3.**
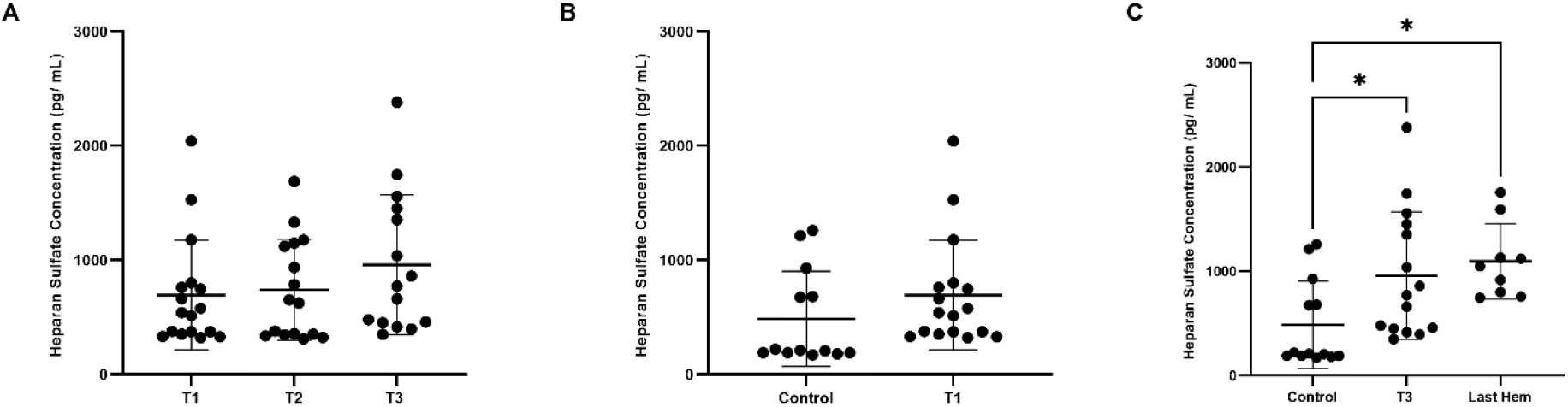
HS shedding in a rat model of lethal hemorrhage shock (65%) and polytrauma plus hemorrhage (40%). All data sets are displayed with individual data points, mean, and standard deviation. Control denotes rats who underwent cannulation, but no trauma. (Fig. 3A) Repeated measures one-way ANOVA analysis of HS levels during accumulating time points denoted T1 (n = 17), T2 (n = 16), T3 (n = 15). (Fig. 3B) Unpaired t-test analysis of Control and T1 plasma HS levels. (Fig. 3C) One-way ANOVA analysis Control (n = 13), T3, and Last Hem (n = 10) plasma HS levels. * p ≤ 0.05. Last Hem denotes plasma collected from the last mL of blood of our rat polytrauma and hemorrhage model. T designates blood collected at various timepoints during our lethal hemorrhage shock model (see methods and Supplemental Figure 1). n = 17 for T1, n = 16 for T2, n = 15 for T3, n = 6 for control, and n = 9 for Last Hem.

### SDC-1 correlation to TM increases with hemorrhage, and is not further elevated by compounded polytrauma

During the course of accumulated lethal hemorrhage, the Spearman coefficient between SDC-1 and TM concentrations, r, steadily rose as blood draw time and % EBV did-r = 0.0311 at T1 (Fig. 4A; p > 0.05), r = 0.401 at T2 (Fig. 4B; p > 0.05), and r = 0.517 at T3 (Fig. 4C; p < 0.05). The Spearman coefficient between SDC-1 and TM concentrations at Last Hem (Fig. 4D) was situated between T2 and T3, calculated as r = 0.312 with p > 0.05. HS correlation to TM and SDC-1 was also measured but did not show consistent or strong trends (See Supplemental Figures 2 and 3).

**Figure 4.**
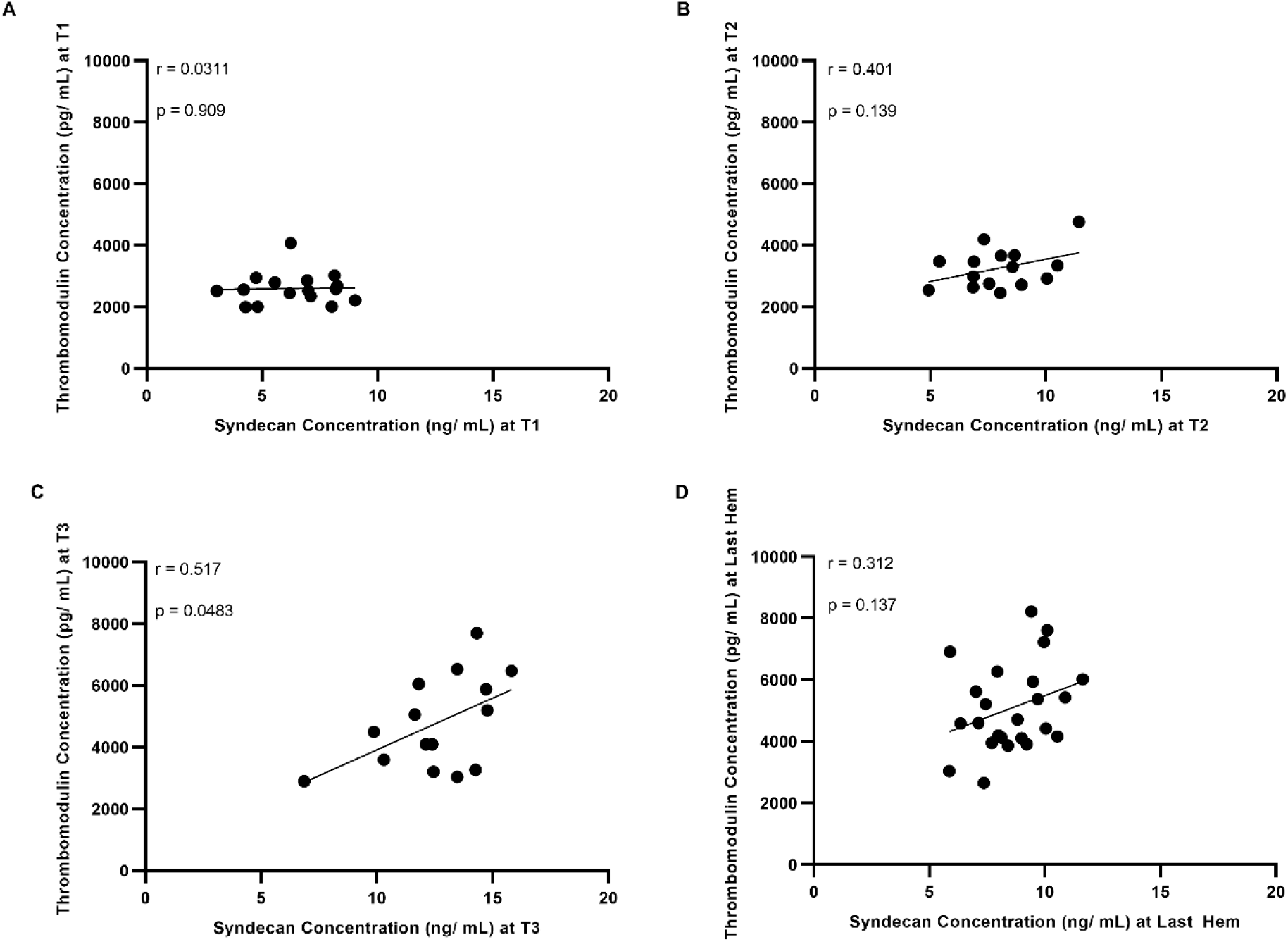
Simple linear regression and correlation analysis of SDC-1 to TM. (Fig. 4A) SDC-1 and TM correlation at time point T1 (n = 15). (Fig. 4B) SDC-1 and TM correlation at time point T2 (n = 15). (Fig. 4C) SDC-1 and TM correlation at time point T3 (n = 15). (Fig. 4D) SDC-1 and TM correlation at Last Hem (n = 16). p ≤ 0.05 is deemed statistically significant.

### Both models significantly elevate lactate and result in a significant drop in PT

Lactate levels in control rats at baseline (denoted “Baseline;” rats who only received cannulation) for subsequent correlation analysis averaged 0.455 mmol/ L ± 0.232 mmol/ L (Fig. 5A). Lactate levels significantly rose to 2.703 mmol/ L ± 0.571 mmol/ L at the end of lethal hemorrhage (denoted “Hemorrhage (T3)”; p < 0.0001) and at the end of polytrauma and 40% hemorrhage (denoted “Last Hem”; 2.655 mmol/ L ± 1.581 mmol/ L p < 0.0001). PT at Baseline averaged 20.66 sec ± 1.465 sec (Fig. 5B).

**Figure 5.**
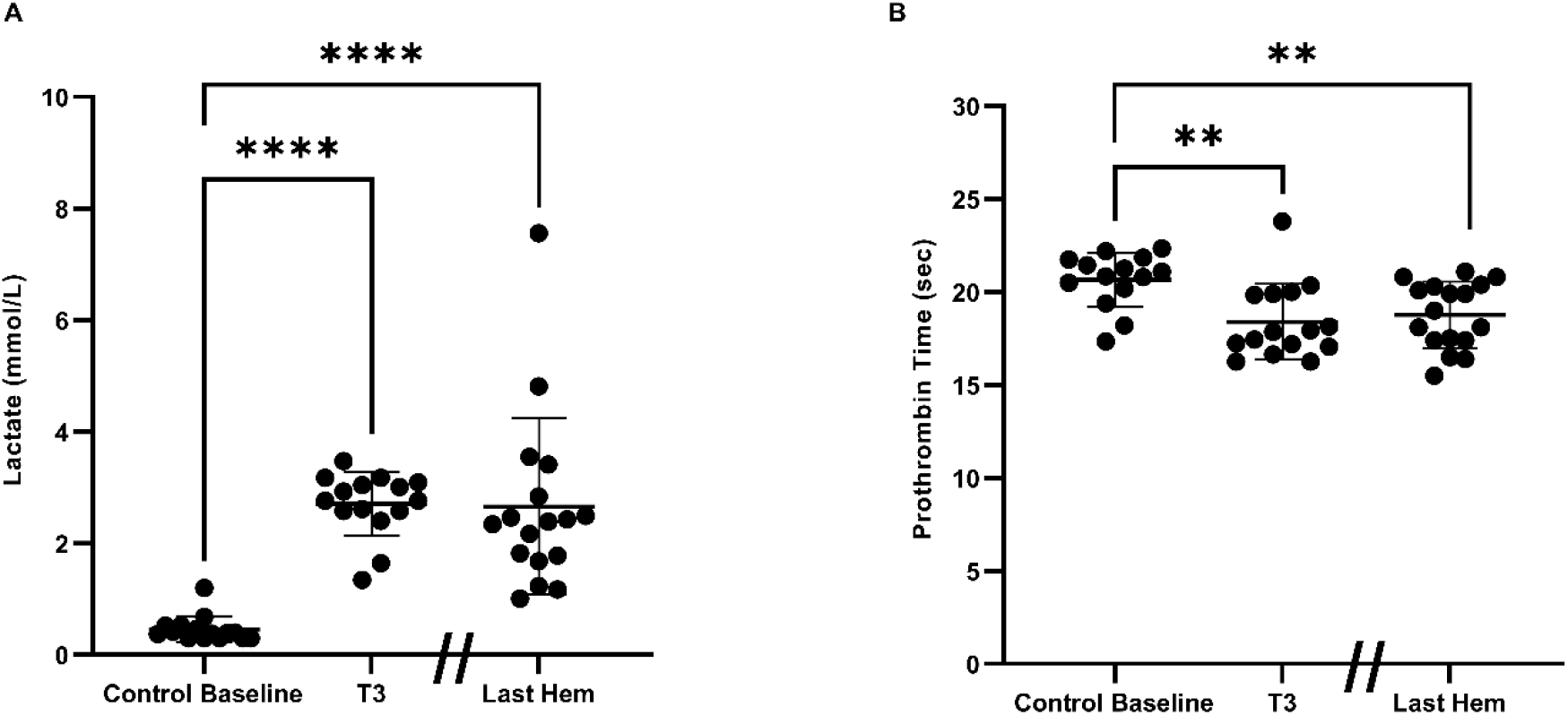
Lethal hemorrhage baseline vs. lactate levels and prothrombin times at the end of both trauma models. All data sets are displayed with individual data points, mean, and standard deviation. (Fig. 5A) Paired t-tests of lactate levels at baseline in control rats that received cannulation only (n = 15; denoted “Baseline”) vs. lactate at the end of 65% lethal hemorrhage (n = 4; denoted “Hemorrhage (T3)”). Unpaired t-tests of lactate levels at baseline in control rats vs. lactate at the end of polytrauma model (n = 17; denoted “Last Hem”). (Fig. 5B) Paired t-tests of prothrombin times (PT) at baseline of control rats (n = 15; denoted “Baseline”) vs. PT at the end of 65% lethal hemorrhage (n = 17; demoted “Hemorrhage (T3)”). Unpaired t-tests of PT levels at baseline of control rats vs. PT at the end of polytrauma model (n = 17; denoted “Last Hem”). ** p ≤ 0.01, **** p ≤ 0.0001.

PT significantly (p < 0.01) decreased to 18.40 sec ± 2.040 sec at Hemorrhage (T3), and at Polytrauma and Hemorrhage end (18.78 sec ± 1.781 sec; p < 0.01).

### SDC-1 tend to positively correlate with lactate and PT, while TM displays the opposite trend

During the time course of accumulating lethal hemorrhage (∆ T3-T1), SDC-1 concentrations tended towards a positive but not significant (p > 0.05) correlation with lactate and PT (Spearman coefficients of 0.148 and 0.319; Fig. 6A and 6C, respectively). Likewise, at the Last Hem blood draw, SDC-1 also tended (p > 0.05) to correlate positively with lactate and PT (Spearman coefficients of 0.465 and 0.437; Fig. 6E and 6G, respectively). TM, on the other hand, while still not significant (p > 0.05), tended to negatively correlate with lactate and PT (Spearman coefficients of -0.203 and -0.124; Fig. 6B and 6D, respectively). Similarly, at the Last Hem blood draw, TM tended (p > 0.05) to correlate negatively with lactate and PT (Spearman coefficients of -0.213 and -0.0774; Fig. 6F and 6H, respectively).

**Figure 6.**
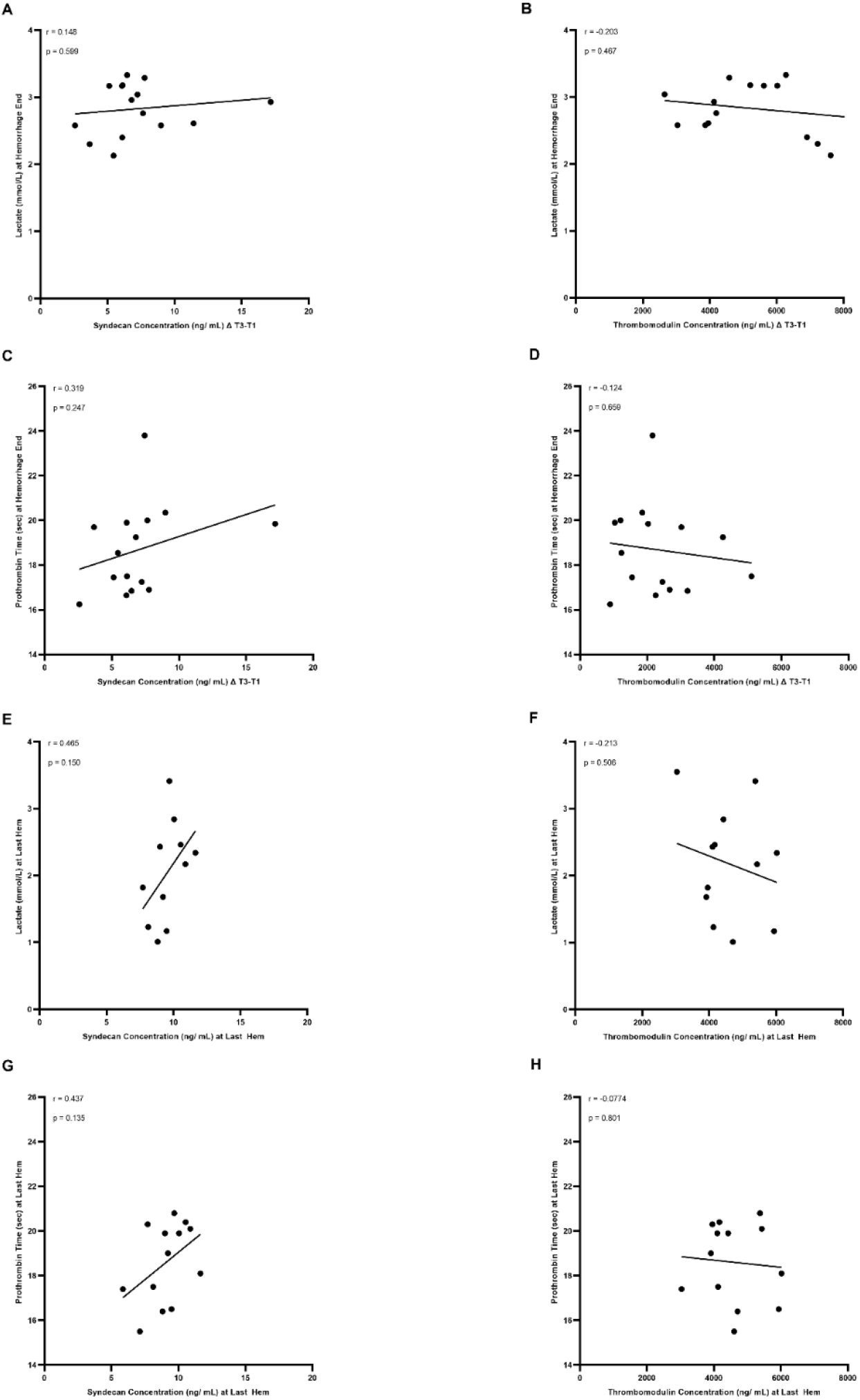
Simple linear regression and correlation analysis of the change in T3 to T1 (denoted ∆T3-T1) of SDC-1 and TM concentrations correlated to lactated levels and PT. (Fig. 6A) SDC-1 correlation to lactate at ∆ T3-T1 (n = 15). (Fig. 6B) TM correlation to lactate at ∆ T3-T1 (n = 14). (Fig. 6C) SDC-1 correlation to PT at ∆ T3-T1 (n = 15). (Fig. 4D) TM correlation to PT at ∆ T3-T1 (n = 15). (Fig. 6E) SDC-1 correlation to lactate at Last Hem (n = 11). (Fig. 6F) TM correlation to lactate at Last Hem (n = 12). (Fig. 6G) SDC-1 correlation to PT at Last Hem (n = 13). (Fig. 6H) TM correlation to PT at Last Hem (n = 13). p ≤ 0.05 is deemed statistically significant.

## Discussion

In this study, we observed the change in plasma levels of three common markers typically associated with endotheliopathy-SDC-1, TM, and HS, which are believed to be associated with the increased breakdown and shedding of the glycocalyx (6, 8) followed by hemorrhage or polytrauma with hemorrhage. While we delineated a significant, acute elevation in end plasma levels of SDC-1 (Fig. 1C), TM (Fig. 2C), and HS (Fig. 3C), we also observed how these biomarkers of endotheliopathy shed as a functional correlation to time and volume of hemorrhage, trauma, lactate, and coagulopathy.

Both SDC-1 and TM plasma levels continued to significantly rise over the course of lethal hemorrhage (Fig. 1A and 2A, respectively) and were detected as early as at 4 and 6 min following 20-30% hemorrhage (Fig. 1B and 2B, respectively). Plasma levels of HS, on the other hand, did not significantly change within the similar timeframe of observation (Fig. 3A-B). The presence of increased hemorrhage in the 65% lethal hemorrhage model generated a significant increase in end plasma SDC-1 levels (at 16-19 min) compared to the polytrauma and 40% hemorrhage model (at 24 min on average) (Fig. 1C), suggesting that SDC-1 shedding might be more dependent on total hemorrhage volume or the speed of blood loss rather than the presence of additional trauma. End lethal hemorrhage volume and the presence of additional trauma did not significantly impact end TM and HS plasma levels (Fig. 2C and 3C, respectively), however, suggesting that with decreased hemorrhage amount, the addition of compounded trauma is adequate to increase TM and HS shedding to that of lethal hemorrhage levels. Certainly, the polytrauma conducted in this model was severe but not the major source of bleeding that led to hemorrhagic shock, which is consistent with fact that hemorrhage is a leading cause of preventable injury-related death during prehospital phase of trauma.

Agonists of TM, HS, and SDC-1 shedding in vivo under certain physiological conditions include growth factors, chemokines, stress-related agonists, sheddases/ matrix metalloproteinases, and even extracellular vesicles (9-13). Additionally, hypotension during and following hemorrhage, and subsequently lowered shear rate and shear stress to below physiological levels, can result in glycocalyx shedding and shedding of the aforementioned markers (14). The addition of polytrauma on top of hemorrhage might activate different shedding mechanisms that account for similar plasma levels of TM and HS at both a high hemorrhage and low hemorrhage with compounded trauma, while increased hemorrhage (and thus alterations in vessel tone and shear stress) might further contribute more to enhanced plasma levels of SDC-1 than additional trauma. Previous studies in our lab (data not shown) have found that pro-inflammatory and anti-inflammatory cytokines, including the interleukin family, vascular endothelial growth factor, interferon gamma, and tumor necrosis factor alpha, are undetectable in acute samplings of trauma blood (T1, T2, T3, and Last Hem). These cytokines become apparent, however, in the later hours of the polytrauma and hemorrhage model (15, 16).

During the course of accumulating 65% lethal hemorrhage, the Spearman coefficient between SDC-1 and TM steadily rose but did not reach significance until the T3 time points (Fig. 4A-C). At the end of the polytrauma and 40% hemorrhage model, the Spearman coefficient between SDC-1 and TM reflected similarly to that between T2 and T3 time points but was not equally significant (Fig. 4D). The ability for SDC-1 and TM to positively and significantly correlate with each other is not surprising, considering that the two markers of endotheliopathy have been known to have similar correlation statistics in other markers of trauma, including COVID-19, myocardial infarction complicated by cardiogenic shock, and sepsis (17-19). What is novel, however, is detecting a significant correlation within the acute stages of lethal hemorrhage (T3).

Although this study did not provide evidence that the elevation of SDC-1 and TM was directly correlated to the survivability of hemorrhagic shock, the results suggest that an acute countermeasure as early as at the point of injury is required in the treatment of hemorrhagic shock for mitigation of early development of endotheliopathy.

During the course of lethal hemorrhage (∆ T3-T1) and at polytrauma and hemorrhage end, SDC-1 trended towards a positive but not significant correlation with lactate and PT, while TM favored a negative correlation to both (Fig. 6). SDC-1 has been shown to positively and significantly correlate with lactate when measured at ICU admission for sepsis (20), and SDC-1 shedding is associated with hypocoagulability associated with prolonged clot reaction in thromboelastography assays (21-23). These correlation data sets in literature, however, were measured at ICU admission and in human patients, so it is not clear when exactly plasma SDC-1 and TM levels, as well as lactate, were measured. In this study, particularly the polytrauma model, lactate and PT levels were measured at an average of 25.07 min ± 7.404 min, while SDC-1 and TM levels were measured earlier. Therefore, such acute sampling time points might be too early to truly see significant correlation effects between SDC-1 or TM and lactate or PT, and thus we were only able to observe a trend. Consequently, the acute change in lactate or PT may not necessarily be used to predict the degree of endotheliopathy as characterized by the elevation of SDC-1 and TM in acute hemorrhagic shock.

Work characterizing biomarkers of endotheliopathy in our 65% lethal hemorrhage model and polytrauma and 40% hemorrhage model corresponds with prior work that has been done in both rats and in humans (24-30). Both rodent models are relevant to the common patterns of trauma and hemorrhage. We previously identified that rats with 65% hemorrhage did not survive to 60 min after hemorrhage (unpublished data), and that the average time to death was 30 ± 4 min, aligning with the time window of the most casualties on the battlefield (mostly due to non-survivable or decompensated hemorrhagic shock by damage of major blood vessels). The rats in the model of polytrauma and 40% hemorrhage were survivable only for 4 hours, which allowed us to study acute pathophysiologic changes in response to severe polytrauma (visceral injury and extremity trauma) combined with hemorrhagic shock, including ATC, inflammation, and multiple organ failure (31, 32). Endotheliopathy was identified as one of the acute drivers to cause or aggravate ATC, inflammation, and multiple organ failure in various clinical studies (1, 3). However, the mechanism for the onset and development of endotheliopathy associated with trauma and hemorrhage has not been well characterized. Using these two rat models allows us to delineate not only the temporary changes of soluble biomarkers for endotheliopathy associated with trauma and hemorrhage, but also enables us to observe the contributive effects made by severe hemorrhagic shock alone or combined with massive tissue injuries.

This study is not without its limitations. First, although minor, femoral artery and vein cannulation has the potential to contribute to minor soft tissue injury and ischemia within the model, thus not technically making the model a 100% full hemorrhage model (32). Secondly, the T3 sample was pulled at either 16 or 19 min after starting hemorrhage, due to needing one full cc of WB for iSTAT and ST4 measurements, although we did ensure an even distribution of 16 and 19 min blood samples within each T3 statistic.

This is the first paper to characterize and compare the acute effects of a 65% lethal hemorrhage and polytrauma with 40% hemorrhage on vascular endotheliopathy, as well as correlate these findings to acute shock and coagulopathy. Our results are of great relevance to the continued effort towards the identification and characterization of vascular dysfunction, particularly acute endotheliopathy, for early interventions in combat casualty care, as we have demonstrated that not only is there a significant temporal elevation in early SDC-1 and TM, but that there might also be a trend for these markers to acutely predict downstream consequences of these trauma indications.

## Methods

### Fixed-Volume Hemorrhage Model

Male Sprague-Dawley rats (375–425 g) were anesthetized with 1.5% to 2% isoflurane/100% oxygen through a nose cone and allowed to breathe spontaneously. Cannulas were placed in the left femoral artery and vein for monitoring arterial blood pressure and for withdrawing blood and injection. Rats were then bled in a controlled manner through the arterial and venous cannula to a total EBV of 65% occurring within 20 minutes. EBV was calculated as 0.06 x Body Weight + 0.77 (33). Whole blood (WB) was drawn for fixed volume hemorrhage in a fast-to-slow pattern, where an initial 40% of total amount of hemorrhage was removed in the first two minutes through femoral artery, followed by 1.5 ml every 2 minutes in the next 8 minutes, and then 1 ml every 3 minutes till the total hemorrhage amount was achieved. The shed blood was collected in sodium citrate (1:9 parts citrate:WB) from rats undergoing hemorrhage at time-points (T) designated T1 (pooled shed blood collected at 4 and 6 min), T2 (pooled shed blood collected at 8, 10, and 13 min), and T3 (shed blood at 16 or 19 min; data sets pooled) (Fig. 1a). The citrated WB was also collected in cannulated healthy rats as control.

### Polytrauma/Hemorrhage Model

Male Sprague-Dawley rats (350–450 g) were anesthetized with 1.5% to 2% isoflurane/100% oxygen through a nose cone and allowed to breathe spontaneously. Cannulas were placed in the left femoral artery and vein for measurement of arterial blood pressure and to obtain vascular access (blood sampling, resuscitation, and drug administration) respectively. Polytrauma (performed in roughly 10 min) was induced by laparotomy, crush injury to the small intestines, the left and medial liver lobes, and the right leg skeletal muscle and by fracture of the right femur. Hemorrhage immediately followed. EBV was calculated as 0.06 x Body Weight + 0.77. Rats were bled to lower mean arterial pressure (MAP) to 40 mmHg within 5 min and maintained at 40mmHg until 40% of estimated blood volume was removed. The last mL of shed blood (denoted “Last Hem”) was collected subsequent experiments.

### Prothrombin Time

Prothrombin times were assessed on a Stago ST4 Coagulation Analyzer (Stago, Parsippany, NJ) using 50 μL of plasma, following manufacturer protocol.

### Lactate Measurements Using i-STAT

WB was used for i-STAT analysis (i-STAT Handheld; Abbot Laboratories, Chicago, IL) using the CG4+ (03P85-50) test cartridges.

### Plasma Preparation

Plasma from WB was prepared by centrifugation at 5,000xg for 10 min at 4°C.

### Enzyme-Linked Immunosorbent Assays (ELISAs)

ELISAs for SDC-1 (cat. #LS-F12571; LS-Bio; Seattle, WA), TM (cat. #LS-F24220; LS-Bio), and HS (HS; cat. #LS-F74206; LS-Bio) were performed following manufacturer protocol.

### Statistics

Statistical analyses were performed using GraphPad Prism (version 7, GraphPad Software, La Jolla, CA). A resulting p value of less than or equal to 0.05 was considered to be significant. ELISA data sets between T1, T2, and T3 were analyzed using a repeated measures one-way analysis of variance (ANOVA) with a Tukey correction for multiple comparisons. ELISA data sets between Control samples, T3, and Last Hem were analyzed using a one-way analysis of variance (ANOVA) with a Tukey correction for multiple comparisons. Lactate levels and PT at baseline and end of trauma models were analyzed by two-tailed paired t-tests. Correlation data sets were analyzed to find the Pearson r value, p value, and then subsequently analyzed by simple linear regression for trend line fit.

### Study Approval

Research was conducted in compliance with Animal Welfare Act, the implementing Animal Welfare regulations, and the principles of the Guide for the Care and Use of Laboratory Animals. The Institutional Animal Care and Use Committee approved all research conducted in this study. The facility where this research was conducted is fully accredited by the AAALAC International.

## Disclaimer

The views expressed in this article (book, speech, etc.) are those of the author(s) and do not reflect the official policy or position of the U.S. Army Medical Department, Department of the Army, DoD, or the U.S. Government.

## Author Contributions

T.C.C., X.W., designed the study. T.C.C. and X.W. performed the experiments and data acquisition. T.C.C., X.W., D.N.D., and J.A.B. interpreted and analyzed the data. T.C.C. wrote the article. M.A.M, A.P.C., and X.W. provided critical revisions of the article.

D.N.D. and X.W. wrote the animal protocols.

## Acknowledgements

The authors would like to thank Mr. Jeffrey Keesee, Ms. Bin Liu, Dr. Qingwei Zhao, Mr. Stephen McBride, and Ms. Lauren Bonnett for assisting with animal protocol, sample collection, and data collection.

## Tables

**Table 1.** Average accumulated hemorrhage amount at each timepoint of blood draw for 65% Lethal Hemorrhage Model, and average time to completion of 40% estimated blood volume (EBV) of hemorrhage in polytrauma/hemorrhage. In 65% lethal hemorrhage model, blood draws at 4-6 minutes were subsequently binned together as T1 and 8-13 min as T2. T3 designates blood draws at either 16 or 19 min. Average Hemorrhage amounts (EBV%) represent a response to the amount of hemorrhage prior to blood draw. In Polytrauma/40% Hemorrhage Model, “Last Hem” denotes the last 1 ml of shed blood. All data sets represent the sum of all rats used for subsequent data analysis.

| 65% Lethal Hemorrhage Model |  |  |  |
| --- | --- | --- | --- |
| Time of Blood Draw | Average Hem (EBV%) | St.Dev. (EBV%) | n |
| T1 (4 min) | 32.33 | 0.349 | 33 |
| T1 (6 min) | 38.66 | 0.697 | 33 |
| T2 (8 min) | 44.98 | 1.046 | 33 |
| T2 (10 min) | 51.31 | 1.394 | 33 |
| T2 (13 min) | 55.53 | 1.627 | 33 |
| T3 (16 min) | 56.55 | 2.818 | 16 |
| T3 (19 min) | 64.19 | 2.230 | 17 |
| T3 (16 min + 19 min) | 60.48 | 4.608 | 33 |
| Polytrauma and 40% Hemorrhage Model |  |  |  |
|  | Average Time (min) | St.Dev. (min) | n |
| Time of Last Hem Pull | 25.07 | 7.404 | 47 |

